# The effect of susceptible and resistant potato cultivars on bacterial communities in the tuberosphere of potato in soil suppressive or conducive to common scab disease

**DOI:** 10.1101/340257

**Authors:** Jan Kopecky, Zuzana Samkova, Ensyeh Sarikhani, Martina Kyselková, Marek Omelka, Vaclav Kristufek, Jiri Divis, Geneviève G. Grundmann, Yvan Moenne-Loccoz, Marketa Sagova-Mareckova

**Author notes:** Address correspondence to Marketa Sagova-Mareckova,.

## Abstract

Connections between the structure of bacterial communities in suppressive soils and potato resistance to common scab (CS) are not well understood. In this study, one resistant and one susceptible cultivar were grown in a conducive and suppressive field to assess cultivar resistance × soil suppressiveness interactions. The resistant cultivar had a higher Mg content in periderm compare to susceptible cultivar, while suppressive soil had lower pH (5.3 vs 5.9), N, C, P, Ca contents but higher Fe and S compared with the conducive soil. Bacteria and actinobacteria numbers were higher in the conducive soil. Copy numbers of *txtB* gene (coding for a pathogenicity determinant) were similar in both soils but were higher in the conducive soil (for periderm samples) and in the susceptible cultivar (for conducive soil samples). Taxonomic microarray analysis and Illumina sequencing of 16S rRNA genes amplicon showed that bacterial community differed between resistant vs susceptible cultivar and to a lesser extend between suppressive vs conducive soil. Bacteria participating in soil suppression belonged to *Pseudomonadaceae, Bradyrhizobiaceae, Acetobacteraceae* and *Paenibacillaceae*, while resistant cultivars selected a bacterial community resembling that of the suppressive soil, which was enriched in *Nitrospirae* and *Acidobacteria*. Thus, the analysis of soil suppressiveness×cultivar resistance interactions enabled to gain new insight to CS control in the field.

**IMPORTANCE:** It was demonstrated that potato cultivars susceptible and resistant to common scab select differing bacterial community and above that this trait is further modified in suppressive and conducive soil. Common scab severity was diminished by either resistant cultivar or suppressive soil but without additive effect between them. Out of the two factors, potato cultivar had a more significant influence on tuberosphere bacterial community composition than soil. Results highlighted the usefulness of both cultivar resistance and soil suppressiveness traits in understanding and managing disease control of crops.

---

Suppressive soils were described as soils in which disease severity remains low, in spite of the presence of a pathogen, a susceptible host, and climatic conditions favorable for disease development (1, 2). Relatively few soils with suppressive character have been described in the world to date (3). They represent a unique model for studying plant disease control. It is of prime interest to conserve their functioning, and ultimately, these soils may help us to learn how to establish suppressive character of soil at other sites (4).

It is agreed that soil suppressiveness is related to soil health and crop productivity through physico-chemical conditions or microbial communities (5, 6). Yet, suppressiveness is difficult to understand because in soil, (i) most physico-chemical factors are not independent from one another, (ii) phytopathogenic taxa may display genetic and functional diversity, and (iii) microbial communities are composed of many taxonomic groups with unknown functions and their structure is affected not only by soil but also by the plant host (4). Therefore, the assessment of suppressive soils may necessitate to target potential interactions between all these ecological factors.

Common scab (CS) of potatoes is a soil-borne disease caused by *Streptomyces* spp. that produce thaxtomin phytotoxins, and for which suppressive soils were reported in the USA (6, 7). In these systems, disease control is largely attributed to biological interactions (mostly competition and antagonism) between plant-beneficial microbiota and pathogens mediated via antibiotic production or enzymatic activities (4, 8). In one situation, nonpathogenic *Streptomyces* spp. were correlated with CS suppressiveness (6), and it was also hypothesized that other actinobacteria may be involved in this disease suppression (4).

In our previous investigations, two sites (Vyklantice and Zdirec) from the Czech Republic, where CS suppressive and conducive fields occur next to each other were studied both in field trials (9, 10) and pot experiments (11). The CS suppressive character of the fields differed according to field location, presumably in relationship to local soil chemical properties (9). Potato cultivars susceptible and resistant to CS have different ecophysiologies, and they differ in chemical composition of the potato periderm, which may further influence microbial community structure (12). Therefore, our hypothesis was that pathogen control via plant-enriched taxa occurring in the tuberosphere was a trait associated with both the suppressive soil and cultivar.

CS-susceptible and resistant potato cultivars have not been compared yet in terms of their respective interactions with the soil microbial community, so that was the focus of the current work. We wanted to assess the relative importance of suppressive soil and resistant cultivar as ecological factors shaping bacterial community properties in CS disease suppression conditions, and identify the corresponding bacterial taxa. We used a field set-up that included a combination of both disease suppressive vs conducive soils, and resistant vs susceptible cultivars. Bacterial community structure in soil and potato tuberosphere was assessed by 16S rRNA taxonomic microarray and Illumina sequencing, and these results were compared to CS severity observed on tuber surface, quantity of thaxtomin biosynthetic genes *txtB*, quantities of total bacteria and actinobacteria and soil and periderm chemical characteristics.

Currently, next generation sequencing is the preferred approach to study bacterial communities. The method enables high throughput and financially-efficient description of bacterial taxa present in a sample. However, the method has its drawbacks. In particular, it is not highly quantitative so only a relative quantity of determined taxa can be assessed, and also it may be difficult to separate sequencing errors from real diversity (13). In comparison, taxonomic microarrays as a method for bacterial community assessments are more laborious (so are slower) and give results only for a selected group of taxa. However, these taxa can be purposefully selected and aimed at determining particular characteristics of bacterial community, and microarrays are semi quantitative (14), which makes microarrays a useful complement to Illumina sequencing. In this work, we used a validated taxonomic microarray focusing on disease suppressive soils and bacterial taxa possessing plant growth-promoting and antagonistic traits in soil environments (15, 16), which was further extended with probes focusing on CS pathogens. The microarray approach was combined with amplicon Illumina sequencing to obtain deeper insight to relationships of bacterial communities in cultivar resistance and soil suppressiveness.

## RESULTS

### Common scab severity and quantities of thaxtomin biosynthetic genes

In conducive soil H, severity of CS (resulting from natural field infestation) was significantly higher in susceptible cultivar Agria than resistant cultivar Kariera (Fig. 1; ANOVA, p<0.001). In suppressive soil L, CS severity did not differ between the cultivars, and was as low as for the resistant cultivar in conducive soil. The number of *txtB* gene copies was similar in both soils (Supplementary Tables S2A and S4A), while in periderm it was significantly higher (p=0.006) in conducive than suppressive soil (Supplementary Tables S2B and S4B). The two cultivars grown in the same soil had comparable quantities of *txtB* gene copies in their periderm, yet, the number of *txtB* gene copies was significantly higher for susceptible cultivar Agria in conducive soil than in suppressive soil. In summary, CS control required resistant cultivar (independently of the soil) or suppressive soil (for susceptible cultivar).

**FIG 1.**
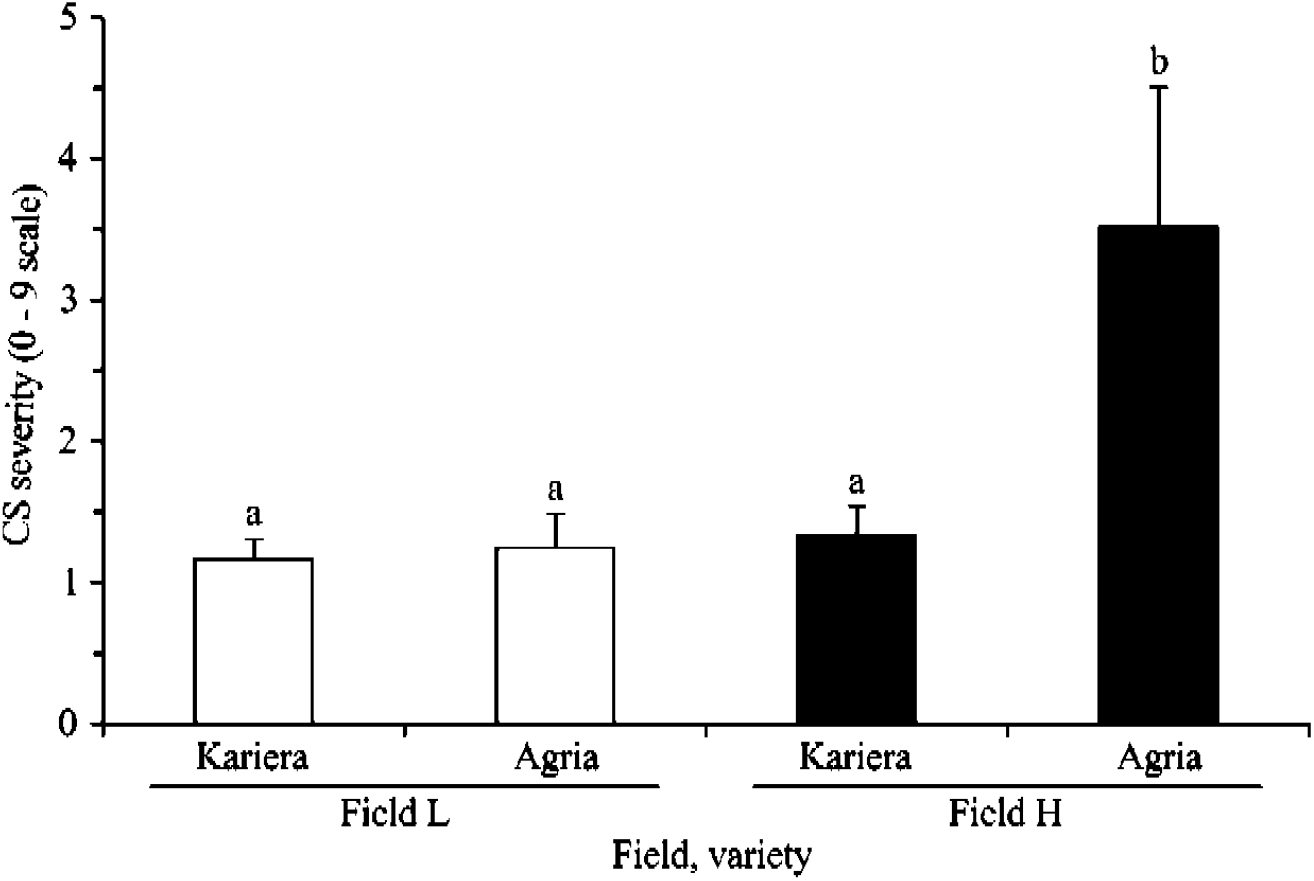
Severity of common scab of susceptible cultivar Agria and resistant cultivar Kariera in suppressive and conducive soils (means ± standard deviations, n = 4). Statistical significance between treatments (ANOVA) are shown with letters a and b.

### Chemical composition of tuberosphere soil and periderm

In tuberosphere, contents of N, C, P, Ca, Fe and soil pH were significantly higher in conducive than suppressive soil (ANOVA; all p<0.001), while S content was significantly higher in suppressive soil (ANOVA; p<0.001). Ca content was significantly higher in bulk soil than in tuberosphere of both soils (ANOVA; p<0.001; Supplementary Tables S3A and S4A). In periderm, N content was significantly higher in both cultivars from suppressive soil (ANOVA; p<0.001), Ca content was significantly higher in susceptible cultivar Agria in both soils (ANOVA; p=0.011), and Mg content was significantly higher in resistant cultivar Kariera in both soils (ANOVA; p<0.001). Fe content was significantly higher in conducive soil in both cultivars (ANOVA; p=0.035) and it was higher in the resistant cultivar in both soils (ANOVA; p=0.006; Supplementary Tables S3B and S4B).

In summary, CS control was connected either with lower content of N, C, P, Ca, Fe and lower soil pH in tuberosphere in suppressive soil, or with higher content of Mg in periderm of the resistant cultivar. In addition, S content was significantly higher in tuberosphere for the combination of suppressive soil x resistant cultivar.

### Quantities of total bacteria and actinobacteria

In tuberosphere, the quantities of bacteria (ANOVA; p<0.001) and actinobacteria (p=0.006) were higher in conducive than in suppressive soil. In suppressive soil the quantity of both bacteria (p=0.011) and actinobacteria (p=0.019) was significantly lower in plant tuberosphere compared to bulk soil (Supplementary Tables S2A and S4A). In periderm, the quantity of actinobacteria (ANOVA; p=0.021) was significantly higher in conducive than in resistant cultivar Kariera suppressive soil in both cultivars and was also significantly higher in susceptible cultivar Agria than Kariera in conducive soil (Supplementary Tables S2B and S4B). In summary, quantities of total bacteria and actinobacteria depended on soil (suppressive vs conducive) × cultivar (resistant vs susceptible) × compartment (periderm vs tuberosphere vs bulk soil) combination, with a trend for lower number(s) in supressive soil and resistant cultivar.

### Bacterial community composition in bulk soil and tuberosphere by microarray analysis

The 16S rRNA taxonomic microarray previously validated for bacterial community analysis of rhizosphere soil samples (15, 17) was expanded for coverage of the genus *Streptomyces*, including pathogen species *S. scabies* and relatives (Supplementary Table S1).

Non-metric multidimensional scaling (NMDS) plot of sample distances calculated from microarray data demonstrated that bacterial communities in conducive and suppressive soils were distinct, and in tuberosphere they were also influenced by cultivar (Fig. 2A). According to PERMANOVA, cultivar explained 42% variability and field site 13% variability. In particular, bacterial community in tuberosphere of the susceptible cultivar was separated from those of the resistant cultivar and bulk soil. Bacterial communities were significantly closer to each other within conducive or suppressive soil when compared to all samples (PERMANOVA; p=0.003) and samples of bacterial communities were significantly closer within each cultivar (PERMANOVA; p<0.001) but not within each bulk soil. In tuberosphere, bacterial communities of resistant cultivar Kariera differed between the soils (PERMANOVA; p<0.001), while bacterial communities of susceptible cultivar Agria did not differ significantly between the two soils but differed from those of resistant cultivar Kariera in each soil (PERMANOVA; p=0.029).

**FIG 2.**
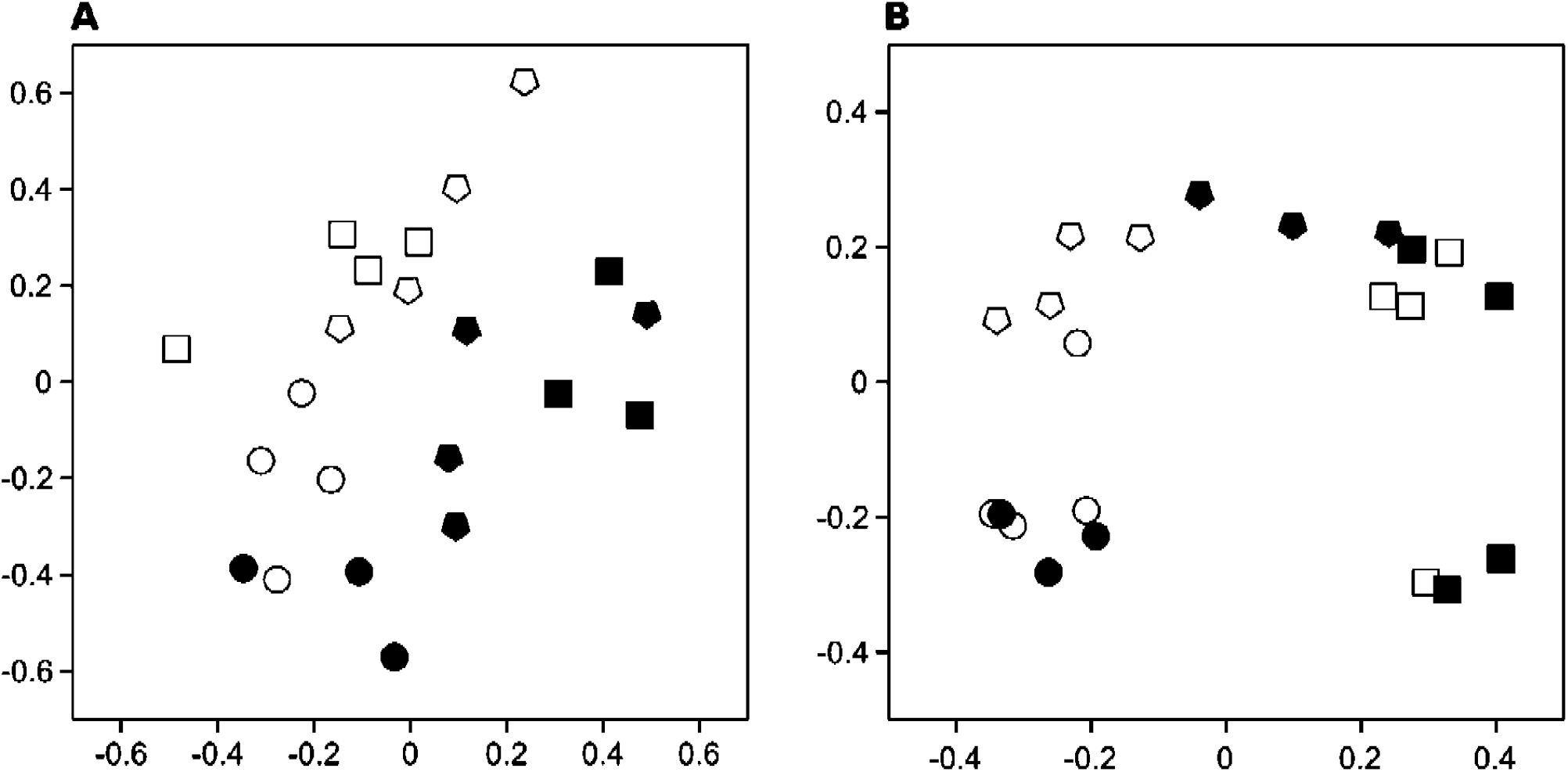
Bacterial communities in tuberosphere of CS-susceptible cultivar Agria (circles) and CS-resistant cultivar Kariera (squares), and in bulk soil (pentagons) assayed by 16S rRNA taxonomic microarray (A) and 16S rRNA gene Illumina amplicon sequencing (B) in CS-suppressive (open symbols) and CS-conducive soils (grey symbols). Non-metric multidimensional scaling of distance matrices was based on Yue-Clayton theta calculator.

In summary, our 16S rRNA probe set was expanded to target also the CS pathogens and other *Streptomyces* taxa. The resulting taxonomic microarray evidenced significant differences in bacterial community features when comparing suppressive vs conducive soil and resistant vs susceptible cultivar.

### Discriminant microarray probes according to soil and potato cultivar

Microarray probes contributing to separation of suppressive from conducive soil were Brady4 (targeting the family *Bradyrhizobiaceae*), Pseu33 (*Pseudomonadaceae*), Aceto3A (*Acetobacteraceae*), and PalgiG3 (*Paenibacillaceae*), which were significantly higher in suppressive soil, as well as Janaga 2 and 3 (targeting the genus *Janthinobacterium*), which were significantly higher in conducive soil (Supplementary Table S5A). These differences seemed enhanced by tuberosphere of susceptible cultivar Agria, based on lower hybridization of probes targeting the genus *Streptomyces* or families *Rhizobiaceae* or *Bradyrhizobiaceae* but higher signals for probes aiming at pathogenic *Streptomyces scabiei* group.

Probes contributing by their higher signal intensity to separation of resistant cultivar and bulk soil from the susceptible cultivar Agria were Rhizo157 and Rzbc1247 (both targeting the family *Rhizobiaceae*), Strepto1, 2, and 3 (targeting the genus *Streptomyces*) and Plancto4.mB (targeting the phylum *Planctomycetes*) (Supplementary Table S5B). Relatively small numbers of probes discriminated between the two soils when assessing bulk soil samples (only 2 probes), tuberospheres of the resistant cultivar Kariera (13 probes), and tuberospheres of the susceptible cultivar Agria (26 probes). Differences between samples from a same soil implicated somewhat comparable numbers of discriminating probes (Fig. 3A).

**FIG 3.**
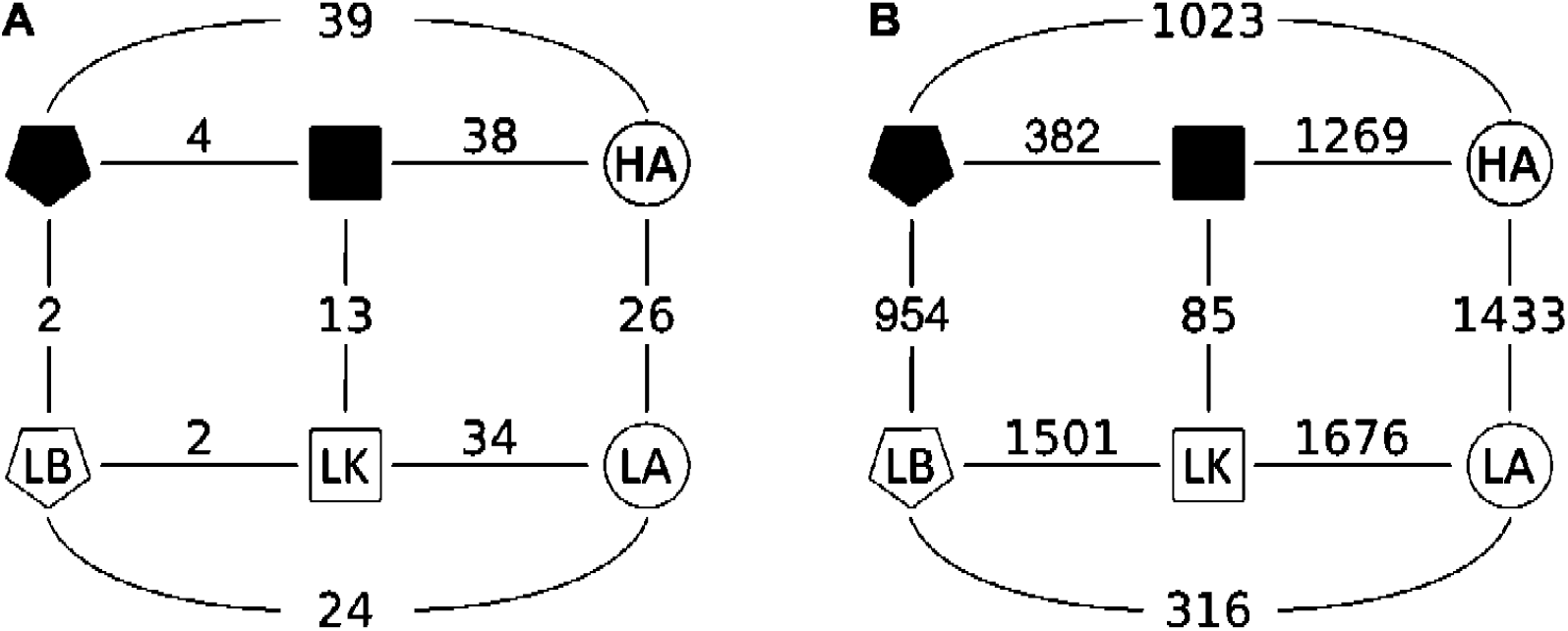
Differences between bacterial communities in tuberosphere of CS-susceptible cultivar Agria (circles) and CS-resistant cultivar Kariera (squares), and in bulk soil (pentagons) assayed by 16S rRNA taxonomic microarray (A) and 16S rRNA gene Illumina amplicon sequencing (B) in CS-suppressive (L, open symbols) and CS-conducive (H, grey symbols) soils. Numbers indicate the numbers of probes (A) and OTUs (B) significantly contributing to the difference between samples in pairwise comparisons (Metastats, p < 0.05).

In summary, six probes targeting contrasted bacterial taxa discriminated between suppressive and conducive soils. Higher probe signals for CS pathogen were found with the susceptible cultivar.

### Bacterial community composition in bulk soil and tuberosphere by Illumina sequencing

A total of 1,213,004 16S rRNA gene sequence reads were obtained, out of which 944,597 (i.e. 78%) were mapped to 4001 OTUs. On a NMDS plot, bacterial communities of resistant cultivar Kariera, susceptible cultivar Agria and bulk soil were separated from one another within each field (Fig. 2B). When comparing both fields using significantly different OTUs (Metastats p<0.05), bacterial communities of bulk soils were separated, whereas they overlapped when comparisons were made for cultivar Kariera, or for cultivar Agria. The number of discriminating OTUs (Fig. 3B) was only 85 between both fields for resistant cultivar Kariera, 382 between bulk soil and resistant cultivar Kariera in conducive soil, and 316 between bulk soil and susceptible cultivar Agria in suppressive soil, whereas the other pairwise differences between treatments implicated 954-1676 discriminating OTUs.

The relative proportion of bacterial phyla did not differ significantly between bulk soils, except that *Actinobacteria* were higher and *Acidobacteria* lower in suppressive than in conducive soil (Supplementary Fig. S1A). Based on comparison with bulk soil, the tuberosphere communities implicated (i) an increase in relative proportion of *Chloroflexi* and decrease in that of *Verrucomicrobia, Gemmatimonadetes, Planctomycetes* and *Proteobacteria* in both cultivars (in the two fields; Supplementary Fig. S1A), (ii) an increase in relative proportion of *Bacteroidetes* (particularly the family *Sphingobacteraceae*) in resistant cultivar Kariera (in the two fields; Supplementary Fig. S1D), and (iii) an increase in relative proportion of *Firmicutes* (especially the family *Paenibacillaceae*) and *Actinobacteria* (especially the orders *Gaiellales, Micrococcales, Frankiales* and *Streptomycetales*) in susceptible cultivar Agria (in the two fields; Supplementary Fig. S1B,C). This increase in *Streptomycetales* was contributed by OTU 176, to which also the CS pathogen belongs. However, other members of this OTU contributed more significantly because this OTU was defined by centroid sequence, which was at 2.1-2.7% distance from the pathogen (Supplementary Table S6B).

Diversity of bacterial communities based on rarefaction curves showed that in both soils the lowest diversity was in tuberosphere of susceptible cultivar Agria. Above that, the curves were mixed for bulk soil and both cultivars in suppressive soil, while in conducive soil they were mixed only for bulk soil and tuberosphere of resistant cultivar Kariera. Therefore, the diversity in bulk soil was more heterogeneous in suppressive soil and differed more between cultivars in conducive soil (Supplementary Fig. S2).

In summary, Illumina sequencing of 16S rRNA genes evidenced differences between resistant and susceptible cultivars, the latter displaying lower bacterial diversity. Differences were also found and between suppressive and conducive soils, but mainly for bulk soil samples.

### Discriminant OTUs according to soil and potato cultivar

When considering bulk as well as tuberosphere soil samples based on the discriminating OTUs (Metastats, p<0.05), suppressive soil was enriched in *Plantomycetes* (OTUs 385, 2780) and *Bacteroidetes* (OTUs 1402, 1154, 1408) and conducive soil in *Actinobacteria* (OTUs 355, 1230, 886) and *Chloroflexi* (OTU 1478). Different OTUs separating the two soils were found for *Proteobacteria* (with OTUs 92, 253, 68, 592, 835 enriched in suppressive soil vs OTUs 369, 899, 2391, 1832, 2001 in conducive soil) and *Firmicutes* (OTU 3391 enriched in suppressive soil vs OTUs 2120, 2105, 1772 in conducive soil) (Supplementary Table S6A).

The tuberosphere of Agria was enriched in taxa from orders *Frankiales* (*Frankiaceae, Acidothermaceae, Geodermatophilaceae*; OTUs 63, 20, 54, 117) and *Micrococcales* (*Intrasporangiaceae*; OTUs 13, 10) (Supplementary Table S6B, Supplementray Fig. S1B) and class *Gemmatimonadetes* (*Gemmatimonadaceae*; OTU 36), while tuberosphere of Kariera displayed significant enrichment in taxa from phylum *Acidobacteria* (OTUs 51, 275, 143, 138, 76; Supplementary Table S6B, Supplementray Fig. S1A). Tuberosphere communities of both cultivars were also separated by different OTUs belonging to the same taxonomic groups. These discriminating taxa included (i) *Betaproteobacteria* i.e. *Burkholderiales* (OTU 38 in Agria vs OTU 199 in Kariera), (ii) *Alphaproteobacteria* i.e. *Sphingomonadales* (OTU 1 in Agria vs OTUs 48, 696, 282 in Kariera), (iii) *Actinobacteria* i.e. *Propionibacteriales* (OTU 138 in Agria vs OTU 6 in Kariera), *Gaiellales* (OTUs 21, 140, 41, 114, 104, 19, 8, 309, 213, 110, 23, 946 in Agria vs OTUs 16, 12, 30, 46, 107, 105, 24 in Kariera) and *Solirubrobacterales* (OTU 31 in Agria vs OTU 69 in Kariera), and (iv) *Chloroflexi* (OTUs 18, 26, 77, 137 in Agria vs OTUs 4, 55, 164, 29, 123, 284, 74 in Kariera) (Supplementary Table S6B).

In summary, suppressive soil was enriched in *Plantomycetes* and *Bacteroidetes* and conducive soil in *Actinobacteria* and *Chloroflexi*, and soils also differed in their *Proteobacteria* and *Firmicutes* profiles. Resistant and susceptible cultivars differed based on 1 *Gemmatimonadetes*, 5 *Acidobacteria*, 6 *Proteobacteria*, 29 *Actinobacteria* and 11 *Chloroflexi* discriminant OTUs.

## DISCUSSION

Disease suppressiveness of Vyklantice soil L was shown in previous studies (9, 11) and since the soil has been regularly interrupted by cropping sequence over long time it can be considered naturally suppressive to the disease (2). In the current experiment, CS disease severity was assessed again in suppressive soil L and conducive soil H, under field conditions, but this time the significance of cultivar status was also investigated, by comparing two potato cultivars susceptible or resistant to CS. Results indicated that CS severity of the susceptible cultivar grown in the suppressive soil was as low as for (i) the resistant cultivar in the same soil, and (ii) the resistant cultivar in the conducive soil. These results showed the effect of soil suppressiveness and cultivar resistance on disease severity, and in particular the similar potential of both types of control mechanisms.

In contrast to our previous work, the current set-up enabled to compare soil suppressiveness and host resistance, and we further identified that soil suppressiveness was significantly related to (i) low quantities of bacteria and actinobacteria and (ii) low C, N contents and high S content in the tuberosphere, while cultivar resistance was related to high Mg content in the periderm. In spite of extensive literature on the positive effect of S on CS severity (reviewed in (18)) it does not seem that S was a key factor for CS control in our soils. A combination of both cultivar resistance and soil suppressivness was related to (i) increased Fe content in both tuberosphere and periderm, and (ii) changes in pathogen populations, which is in agreement with our previous work (11), in which additions of available Fe to soil decreased CS severity.

Out of the parameters related to CS severity, the quantity of bacteria including actinobacteria was previously linked with the amount of available carbon, which differed between conducive and suppressive soils (e.g. (19)). Specifically, the quantity and community structure of actinobacteria were connected to pathogen population and its interactions (11, 20). Further to that the quantity of pathogenic streptomycetes (based on numbers of *txtB* genes) did not change with soil suppressiveness or cultivar in tuberosphere and bulk soil, but in suppressive soil the number of pathogens decreased in potato periderm possibly due to both microbial interactions and soil chemical conditions depending on location (9, 21). In this study, however, the increased numbers of actinobacteria in periderm of susceptible cultivar did not correspond to pathogenic streptomycetes, so perhaps an antagonistic community of actinobacteria developed there as a response to pathogen infection, similarly as in Rosenzweig et al. (8) or Tomihama et al. (22).

The increase of magnesium in resistant cultivar may be explained by several aspects of CS disease and plant metabolism. Previously, we did not find any correlation between Mg periderm content and CS, probably because the field site effect was more important (12). However, the importance of site Mg concentration was correlated to CS and quantities of thaxtomine gene *txtA* in Lazarovits et al. (23), and similarly Lacey and Wilson (24) found that CS disease severity was related to contents in exchangeable Ca, Mg, and K cations. Generally, Mg has both indirect as well as direct effects on disease (25). Magnesium functions also as a P carrier in plants, and P periderm concentration was negatively correlated with CS in many studies (e.g. (12)). Also, root–microbial activities are key factors that determine plant-available Mg release from soils, and Mg was among nutrients affecting microbial community in potato rhizosphere (26). So, we concluded that Mg connection to resistant cultivar is a result of a combination of several factors, which specifically in our experiment resulted in diminishing of CS severity. Finally, higher Fe content in resistant cultivar than susceptible cultivar regardless of the nutrient content in soil is probably connected to better acquisition of this element from soil, either by the resistant plant itself or through its selected bacterial community as pointed by Sarikhani et al. (11). This possibility was raised also by Inceoğlu et al. (27), where potato cultivars grown in two soils yielded different contents in C, S and P while selecting microbial communities with different functional capabilities

Bacterial community structure was identified as a major aspect of soil suppressiveness to CS (8). In our microarray study, *Pseudomonadaceae, Bradyrhizobiaceae, Acetobacteraceae* and *Paenibacillaceae* were associated with disease suppressiveness, and all four include plant-beneficial species and strains (3, 28). This was not expected, as the last three families were more prevalent in CS conducive than suppressive soil in Michigan (8). Our Illumina sequencing indicated that *Rhizobiales* and *Planctomycetales* were enriched in suppressive soil, as in Cha et al. (29), but *Actinobacteria, Chloroflexi, Bacillaceae*, and *Chitinophagaceae* that increased with CS suppressiveness induced by rice bran (22) were more prevalent in the conducive soil here. Therefore, different biocontrol consortia may be at work in different types of suppressive soils. Several phyla were more prevalent in the susceptible cultivar Agria, whereas others were more prevalent in the resistant cultivar Kariera, including *Nitrospirae* and *Acidobacteria* also enriched in suppressive soil studied by Cha et al. (29).

In this work, Illumina sequencing showed that major effects were due to resistant and susceptible cultivars and microarray analysis also evidenced the effect of suppressive versus conducive soils. The most relevant study to compare is that of Rosenzweig et al. (8) because they also studied CS suppressive soils using next generation sequencing, but they used a single, relatively CS resistant potato variety (Snowden), so they could not demonstrate the effect of cultivar in combination with soil suppressivity. In comparison, Weinert et al. (30) showed the effect of potato cultivar on rhizosphere bacteria selection, while Inceoğlu et al. (27) described more profound influence of soil type over potato cultivar on bacterial rhizosphere communities but studied two soil types not disease suppressive and conducive soil. Therefore, our results brought new insight to bacterial community differences with respect to suppression of CS disease, possibly with a more general implication to other pathogen-plant systems.

Finally, the data obtained by Illumina sequencing and taxonomic microarrays are differently biased and to some extend showed different results. Microarray showed more differences between the two soils, while Illumina sequencing stressed differences between cultivars. In particular, microarray demonstrated increase of *Streptomyces* (*Actinobacteria*), *Bradyrhizobium, Burkholderia* (*Proteobacteria*) or *Nitrospira* (*Nitrospirae*) in suppressive soil, while increase of *Acidibacteria, Pseudomonas, Agrobacterium* and *Janithobacterium* (*Proteobacteria*) in conducive soil. *Streptomyces* and *Rhizobiaceae* were associated with resistant cultivar Kariera, while *Burkholderia* and *Stingomonas* (*Proteobacteria*) were associated with susceptible cultivar Agria. Illumina sequencing discriminated the two soils similarly to microarray by increase of *Bradyrhizobiaceae* and other *Proteobacteria*, but also *Bacteroidetes* and *Firmicutes* in suppressive soil, while different families of *Proteobacteria*, *Actinobacteria* and *Firmicutes* increased in conducive soil. For resistant cultivar Kariera, *Chloroflexi, Gaiellales (Actinobacteria)* were found enriched, while in susceptible cultivar Agria, similarly to microarray results *Burkholderia* and *Sphinomonas* (*Proteobacteria*) and *Actinobacteria* were enriched. We explain the differences by specific characteristics of the two methodological approaches (14–16, 31,32).

In conclusion, we demonstrated the cultivar-specific community selection with respect to their susceptibility or resistance to CS and above that we included comparison of this trait in suppressive and conducive soils. We showed that CS can be controlled either with resistant cultivar or with suppressive soil, with no additive effect between them. Out of the two factors, potato cultivar had a higher effect on tuberosphere bacterial community composition than soil in our experiment. Results highlighted the usefulness of both cultivar resistance and soil suppressiveness traits in understanding and managing disease control of crops.

## MATERIALS AND METHODS

### Sites

Vyklantice is a site where fields suppressive (L for low disease severity), and conducive (H for high disease severity) to potato CS occur at about 100 m distance. The two fields differ in common scab severity by observations over 30 years, while their geological context, soil type, climate and management are similar. The fields were regularly planted under a four-year crop rotation system including rapeseed, clover, potatoes, and grains (wheat or oats) in the past two decades (9).

### Field experiment

Potatoes were planted in the beginning of May and samples of bulk soil, tuberosphere soil and potatoes were collected after 80 days. One susceptible cultivar (Agria) and another resistant (Kariera) to CS were used. Potatoes were all certified seed tubers (common scab below 5% of surface). Four plots of each cultivar were planted at each field and the plots were arranged in a Latin square design. Each plot was planted with 3 rows of 12 potato plants (36 plants) separated by 50 cm of bare soil. Fields were fertilized with 100 kg N/ha (ammonium sulphate, 21% N), 35 kg P/ha (monocalcium phosphate, 35% P_2_O_5_), and 60 kg K/ha (potassium salt, 50% K_2_O). Potatoes were treated with pesticides, once with Nurelle D (EC) (chlorpyrifos, cypermethrin) at 0.6 l/ha to prevent Colorado potato beetle (*Leptinotarsa decemlineata*), and twice with Acrobat MZ (dimethomorph, mancozeb) at 2 kg/ha and Ridomil Gold MZ Pepite (mancozeb, metalaxyl-M) at 2.5 kg/ha against the potato blight. Fungicides were not used.

### Sampling

One potato plant growing in the center of each plot was sampled. Potatoes from this plant were collected and washed in distilled water. All potatoes were carefully pealed using a sterile potato peeler (taking approximately 1 mm thick periderm samples), peels were homogenized and mixed, and subsamples (1 subsample per plant) were taken for further analyses (‘periderm’ samples). Tuberosphere soil samples were collected no further than 3 mm from a potato tuber (for details see (9)). Bulk soil was collected at a distance of approximately 30 cm from the closest plant within each plot using a small sterile spade (1 sample per plot). Common scab severity was evaluated on 20 potato tubers per plot using a 9-degree scale (33). Potatoes used for evaluation were those of the collected plant and several more plants from each plot to achieve at least 20 measurements per plot.

### Soil and potato periderm analyses

To determine total soil C, N, and S contents, 2-g samples of homogenized soil from both bulk soil and tuberosphere were dried, milled, and analyzed using Vario MAX CNS analyzer (Elementar Analysensysteme, Hanau, Germany). To determine all other elements, soil subsamples were leached with boiling nitro-hydrochloric acid (aqua regia) and assessed by optical emission spectroscopy with inductively coupled plasma (ICP-OES) by Aquatest Inc. (Prague, Czech Republic). Analyses of potato periderm were performed by service laboratory of the Institute of Botany (Trebon, Czech Republic). For total nitrogen analysis, 1-3 mg dried periderm was mineralized by modified Kjeldahl method in H_2_SO_4_ with catalyzer at 360°C. For total phosphorus, 20 mg of dried periderm was sequentially decomposed by HNO3 and HClO4. In mineralized samples, both N and P were determined by flow injection analysis with spectrophotometric detection using FIA Lachat QC 8500 analyzer (Lachat Instruments, Hach Company, Loveland, CO). Cation contents in periderm were determined by atomic absorption spectrometry using AAS spectrometer ContrAA 700 (Analytik Jena, Jena, Germany) after mineralization with nitro-hydrochloric acid.

### Soil DNA extraction

Soil samples from tuberosphere and bulk soil were homogenized and subsamples of 0.5 g were used for DNA extraction by method SK described by Sagova-Mareckova et al. (34). Briefly, the method is based on bead-beating and phenol/chloroform extraction followed by purification with CaCl2 and GeneClean Turbo kit (MP Biomedicals, Santa Ana, CA). For DNA extraction from potato periderm, 3 g of periderm samples were fine cut in sterile Petri dish, homogenized, and a 0.3 g subsample was processed in the same way as soil samples to obtain total periderm DNA.

### Real-time PCR (qPCR)

Quantifications were performed with primers eub338f (5’-ACTCCTACGGGAGGCAGCAG-3) (35) and eub518r (5’-ATTACCGCGGCTGCTGG-3’) (36) amplifying a 197 bp fragment of the 16S rRNA gene from bacteria, act235f (5’-CGCGGCCTATCAGCTTGTTG-3’) (37) and eub518r yielding a 280 bp product for *Actinobacteria*, and StrepF (5’-GCAGGACGCTCACCAGGTAGT-3’) and StrepR (5’-ACTTCGACACCGTTGTCCTCAA-3’) yielding a 72 bp amplicon of the thaxtomin biosynthetic gene *txtB* (38), respectively. The analyses were done on a StepOne Plus Real-Time PCR System (Applied Biosystems, Foster City, CA) using 96-well plates with GoTaq qPCR Master Mix (Promega) containing SYBR Green as a double-stranded DNA binding dye. The reaction mixture contained in a total volume of 15 μl: 1× GoTaq qPCR Master Mix, 0.2 μM primers, and 0.2-2 ng diluted DNA sample. For all of the mentioned targets the PCR cycling protocol consisted of initial denaturation at 95°C for 10 min, followed by 45 cycles of 95°C for 15 s, 60°C for 30 s and 72°C for 30 s. Melting curves were recorded to ensure qPCR specificity. Baseline and threshold calculations were performed with the StepOne v. 2.2.2 software. The inhibition was tested by serial DNA dilution from each site, and the dilutions without inhibition of qPCR reactions were used for quantification. All qPCR measurements were done in duplicate. Standards for qPCR were prepared by cloning fragments of target genes from *Streptomyces europaeiscabiei* DSM 41802 in pGEM-T Easy vector system (Promega). After PCR verification and isolation of cloned constructs by Pure Yield Plasmid Miniprep System (Promega), a linear standard was prepared by cleaving with SalI enzyme (New England Biolabs, UK) in a 200 μl reaction mixture containing 1× reaction buffer, 2 μg circular plasmid, and 20 U restriction endonuclease for 2 h in 37°C. The linearized plasmid DNA was purified by phenol-chloroform extraction. Aliquots of linearized and purified standard diluted to 20 ng /μl were stored in −70°C. Results were expressed per g dry soil. All results (including for *txtB*) were above detection limit.

### 16S rRNA gene-based taxonomic microarray

A taxonomic microarray based on DNA probes targeting 16S rRNA genes representing 19 bacterial phyla at different taxonomic levels (15) was used to assess soil samples from potato fields. This microarray was validated previously (15, 17). Twelve probes targeting the genus *Streptomyces*, as well as *S. scabies* and relatives (Supplementary Table S1) were added to the previous probe set (1033 probes) in this study. The probe KO 08 (39) for genus *Streptomyces* was obtained via the *probeBase* server (40) (http://probebase.csb.univie.ac.at). The other 11 probes (20-mers) were designed in this study using ARB sofware (41) (http://www.arb-home.de) and the parameters of the Probe Design function chosen by Sanguin et al. (31, 42). Probe specificity was tested with the Probe Match function in ARB against the reference Silva-104 and with the TestProbe online tool against Silva 126 database (43) (http://www.arb-silva.de), at the weighted mismatch value of 1.5 (15). Hybridization properties of probes (e.g. melting temperature, potential formation of secondary structures and 3’dimers) were further tested *in silico*, according to Sanguin et al. (31, 42).

Universal bacterial primers T7-pA (forward; TAATACGACTCACTATAG-AGAGTTTGATCCTGGCTCAG) and pH (reverse; AAGGAGGTGATCCAGCCGCA) were used to amplify 16S rRNA genes from total DNA extracts (44). Primer T7-pA includes at the 5’ end the sequence of T7 promoter, which enabled T7 RNA polymerase-mediated *in vitro* transcription using purified PCR products as templates. PCR reactions were carried out using Taq Expand High Fidelity (Roche Applied Science, Meylan, France) and cycling conditions described in Kyselková et al. (15). Purified PCR products (50 ng/μl) were fluorescently labelled (Cy3) by *in vitro* transcription, according to Stralis-Pavese et al. (45). Purified RNA was fragmented by incubation with ZnSO_4_, as described (45), and 400 ng subjected to hybridization on the microarray. Each probe was present in four copies per slide, and two slides were hybridized per sample.

Hybridization was carried out according to Sanguin et al. (31). Slides were scanned at 532 nm, images were analyzed with GenePix Pro 7 (Molecular Devices, Sunnyvale, CA), and spot quality was checked visually, as described previously (31). Data filtration was conducted using R 3.3.0 (46) (http://www.r-project.org). Hybridization of a given spot was considered positive when 80% of the spot pixels had intensity higher than the median local background pixel intensity plus twice the standard deviation of the local background. Intensity signals (median of signal minus background) were replaced by their square root value and intensity of each spot was then expressed as a fraction of the total intensity signal of the basic pattern it belongs to (42). Finally, a given feature probe was considered as truly hybridized when (i) hybridization signals were superior to the mean signal of the negative controls and (ii) at least 3 of 4 replicate spots displayed positive hybridization (15).

### Illumina MiSeq sequencing and analysis

From the DNA samples, a fragment of the bacterial 16S rRNA gene including the variable region V4 was amplified by PCR using universal primers with overhang adapters CS1-515F (5’-ACACTGACGACATGGTTCTACAGTGCCAGCMGCCGCGGTAA-3’) and CS2-806R 5’-TACGGTAGCAGAGACTTGGTCTGGACTACHVGGGTWTCTAAT-3’) (47). Construction of amplicon libraries and sequencing using MiSeq sequencer (Illumina, San Diego, CA) were done at the DNA Services Facility, Research Resources Center, University of Illinois (Chicago, MI). Resulting paired sequence reads were merged, filtered, aligned using reference alignment from the Silva database (43), and chimera checked using integrated Vsearch tool (48) according to the MiSeq standard operation procedure (Miseq SOP, February 2018) (49) in Mothur v. 1.39.5 software (50). A taxonomical assignment of sequence libraries was performed in Mothur using the Silva Small Subunit rRNA Database, release 128 (51) adapted for use in Mothur (https://mothur.org/w/images/b/b4/Silva.nr_v128.tgz) as the reference database. Sequences of plastids, mitochondria, and those not classified in the domain Bacteria were discarded. The sequence library was clustered into OTUs using the Uparse pipeline in Usearch v10.0.240 software (13), and the OTU table was further processed using tools implemented in the Mothur software. Distance matrices describing the differences in community composition between individual samples were calculated using the Yue-Clayton theta calculator (52). Analysis of molecular variance (AMOVA) (53) was based on a matrix of Yue-Clayton theta distances. Metastats analysis (54) was used to detect differentially represented OTUs.

### Statistical analyses

Analysis of variance (ANOVA) and Fisher LSD tests were used to test differences between soils and cultivars for soil chemical parameters and copy numbers of bacterial and actinobacterial 16S rRNA genes, and *txtB* genes in soil and periderm samples. Permutational multivariate analysis of variance (PERMANOVA) was used to compare distance matrices between microarray samples (55). AMOVA was used to test differences between distance matrices (Yue-Clayton Theta) between Illumina samples. The distance matrices were plotted by Sammon’s Multidimensional Scaling (56).

### Accession number

MiSeq 16S rRNA gene amplicon sequences have been deposited in the NCBI Sequence Read Archive (www.ncbi.nlm.nih.gov/sra) as BioProject PRJNA474544.

## ACKNOWLEDGEMENTS

This work was supported by the Ministry of Agriculture of the Czech Republic, grants QK1810370 and RO0417. We would like to thank Stefan Green and the DNA Service Facility of the UIC for MiSeq sequencing.

